# Iron homeostasis in the annual and perennial stem zones of *Arabis alpina*

**DOI:** 10.1101/2024.08.13.607737

**Authors:** Anna Sergeeva, Hans-Jörg Mai, Petra Bauer

## Abstract

Plants serve as reservoirs for vital micronutrients, including iron, which they store in bioavailable forms to support growth in subsequent seasons. The perennial life style is preponderant in nature. Annual species allocate iron towards their seeds. However, our understanding of iron homeostasis in perennial plants remains limited. *Arabis alpina* is a perennial model organism. Its perenniating branches undergo secondary growth where they store carbon-rich macromolecules.

In this study, we investigated iron homeostasis in the perennial and annual stem zones (PZ, AZ) of *A. alpina*. We found that both, the wild-type Pajares (Paj) and *perpetual flowering 1* mutant accumulated iron at various developmental stages in the PZ as well as in the AZ. Notably, iron levels in the PZ were found to be approximately two-fold higher than those in the AZ, underscoring the PZ’s enhanced capacity for iron storage, irrespective of flowering status. Iron was predominantly located within plastid-bound ferritin, providing insight into its storage mechanism. Furthermore, gene expression analyses supported the significance of ferritin and demonstrated an enrichment of transcripts related to iron homeostasis within the stems. Distinct patterns of expression among iron homeostasis genes were observed in relation to iron contents in the PZ and AZ, indicating tissue-specific regulatory mechanisms governing iron accumulation.

These findings collectively emphasize the critical function of secondary growth and the PZ as an important site for iron storage in perennial plants, suggesting that future research should further explore the nuances of iron homeostasis signaling in perennial plants.

**Highlight:**

‐ Iron accumulates in the perennial stem zone, and ferritin is a storage form for iron there.
‐ Transcripts of iron homeostasis genes are enriched among genes expressed in the annual and perennial stem zones, yet iron accumulation correlates with different gene expression patterns.

## Introduction

Iron is an important micronutrient for organisms as it is widely used as a redox catalyst and in electron transfer. Although abundant in the Earth crust, iron is largely unavailable to organisms in the oxygenated environment. For plants, this situation is aggravated in alkaline and calcareous soil conditions that are on the rise with an increase of arid and semi-arid regions during climate change (Bontpart et al., 2024). Hence, iron is a precious element, that has to be efficiently mobilized by plants in the soil and preserved in a bioavailable form for the next growth season and generation of plants.

Perennial plants live for longer than one growth season. The perennial life style is wide-spread and several different forms of plant perennialism exist in nature (Friedman, 2020). One common feature of perennial plants is that they store and reuse elements and nutrients rather than pass it on towards the next generation in the form of seeds, so that perennial individuals can sustain unfavorable growth seasons. Wild perennials may have abilities that allow them to thrive on nutrient-poor soils, and these may be associated with nutrient-use efficiency during the transitions between growth seasons. It is of agricultural concern when perennial crops are not sufficiently well adapted to calcareous soil. For example, citrus, peach and apricot production are frequently constrained to regions with calcareous soils. Yet, they frequently suffer from iron deficiency chlorosis on such soils, and to ensure high productivity, each tree is yearly treated with iron fertilizer in high quantities (Abadía et al., 2011; El-Jendoubi et al., 2011). Although iron deficiency responses have been studied in fruit tree seedlings, little is known about the molecular components involved in the iron use capacities of several-year-old trees related to nutrient storage between the growth cycles (Martínez-Cuenca et al., 2021). This appears surprising, but one reason is certainly that molecular studies on iron mobilization and storage are difficult to perform in adult trees.

Many insights into molecular plant physiology stem from studies conducted with genetic plant model systems, most widely used the annual *Arabidopsis thaliana* (Woodward and Bartel, 2018). While *A. thaliana* cannot provide insight into perennial growth habits, a close relative of it does. The polycarpic perennial *Arabis alpina*, Alpine rockcress, is a model system that facilitates understanding the perennial life style (Wötzel et al., 2022). It is a pioneer species often in nutrient-poor areas (Wötzel et al., 2022). Perennial *A. alpina* plants from the reference accession Pajares have a particular shoot architecture in which certain stem segments form flowers and fruits or have elongated lateral branches, and again others have dormant buds (Vayssières et al., 2020). It was found that the perenniating stem segments of *A. alpina* undergo secondary growth and become enriched in carbon storage macromolecules, starch and storage lipids (Sergeeva et al., 2021a; Sergeeva et al., 2021b). The zone in which this occurs has therefore been named “perennial zone” (PZ) (Sergeeva et al., 2021a). The PZ is clearly distinct in its anatomy from the “annual zone” (AZ) where no secondary growth is initiated, carbon storage does not take place but instead flowers and seeds are formed, with the AZ becoming senescent with the ending of the growth season (Sergeeva et al., 2021a). The concrete anatomy and flowering of *A. alpina* is influenced by vernalization through cold periods in a wild type accession from Spain that serves as reference of this species, named Pajares (Paj). Interestingly, a derived flowering time mutant that does not require vernalization, named *perpetual flowering 1* (*pep1-1*) continuously flowers (Albani et al., 2012; Bergonzi et al., 2013). Despite the altered flowering behavior, *pep1-1* stems are divided into a PZ and AZ. Starch and lipid bodies are formed in a tissue-specific pattern in cambium and cambium derivatives in the PZ of *pep1-1*. While starch was confined to the secondary phloem parenchyma, lipid bodies were present in secondary phloem and associated parenchyma, secondary xylem and cambium (Sergeeva et al., 2021a; Sergeeva et al., 2021b). Hence, the perennial trait of nutrient storage in a secondary growth tissue does not depend on vernalization and PEP1. To date, it has not been investigated how iron homeostasis is managed in *A. alpina*. Rarely any studies have addressed the importance of iron allocation and storage in perenniating organs versus senescing ones, as most studies on perennial life aspects focus on the regulation of flowering and bud dormancy control (Zhao and Wang, 2024). While carbon and nitrogen storage were examined in *A. alpina* (Sergeeva et al., 2021a; Sergeeva et al., 2021b), nothing is known about allocation of the micronutrient iron and the tissues involved in in the PZ versus the AZ. It is not known whether any iron homeostasis genes are expressed or differentially expressed between the PZ and AZ or during their formation. Knowing genes and regulatory patterns of iron homeostasis genes important for the perennial life style, might facilitate in the future improving iron use efficiency traits in perennial crops and fruit trees.

Here, we addressed such open questions and we investigated whether the PZ might be specialized in the storage of iron in *A. alpina*. We determined whether iron had accumulated in the PZ or AZ stem segments, whether this occurred in the form of ferritin and in which tissues iron was deposited. Furthermore, we investigated which genes were expressed and regulated in the various stages of secondary growth in the PZ versus the AZ. We found evidence of distinct iron localization patterns in the stems of the perennial model plant and we were able to associate certain patterns of gene expression with iron accumulation in distinct shoot segments. These findings help to understand iron allocation in a perennial species and form the basis for investigating the genetic underpinning and signaling of iron allocation in a perennial plant for future correction of iron deficiency chlorosis through better genetic manipulation of internal iron use efficiency.

## Materials and Methods

### Plant Materials

Plant stem segments and materials used for microscopic and biochemical analysis were collected and described previously (Sergeeva et al., 2021a) (see also Fig. 1A, B).

**Fig. 1.**
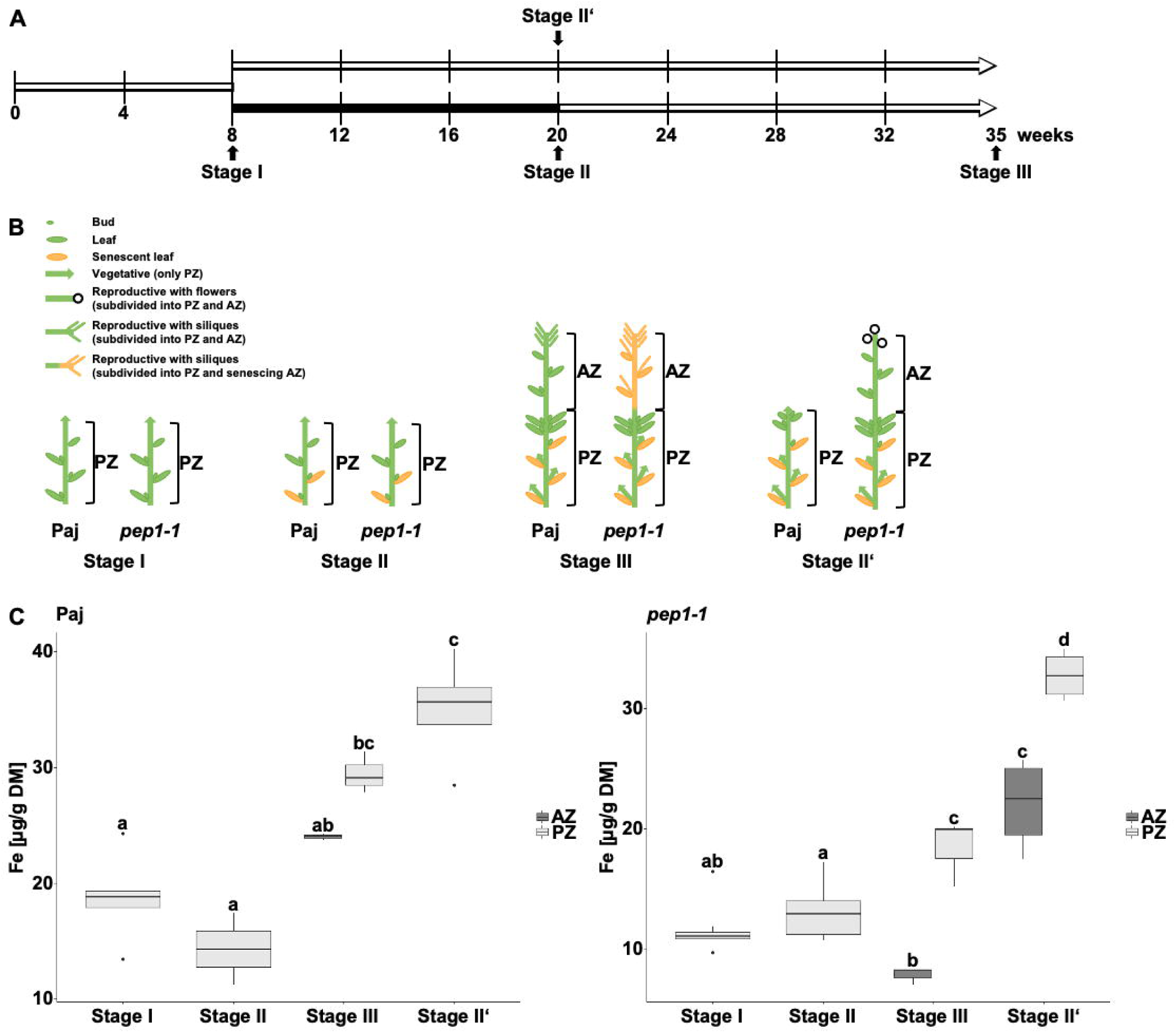
Plant growth scheme and schematic representation of lateral stem stages for *A. alpina* Pajares (Paj, wild type) and its *perpetual flowering 1-1* (*pep1-1*) mutant derivative used in iron content determination. (A) Plant growth scheme and harvesting time points. Open lines, long-day conditions at 20°C; black line, short-day conditions at 4°C (vernalization); four developmental stages, I, II, III and II‘, were harvested. (B) Schematic overview of lateral stem architecture of Paj and *pep1-1* at stages I, II, III and II‘. During harvesting, lateral stems were subdivided into perennial (PZ) and annual (AZ) zones, as specified (Sergeeva et al., 2021a). The schematic lateral stem representation was partially adopted from (Lazaro et al., 2018; Vayssières et al., 2020; Wang et al., 2009). (C) Iron contents per dry matter (DM) in lateral stem internodes in the PZ and AZ at the four developmental stages, as indicated in (A) and (B). Different letters indicate statistically significant differences, determined by one-way ANOVA-Tukey’s HSD test (*P* ≤ 0.05), n = 3-8. The interquartile range (IQR) is indicated by boxes. The median of the data is represented by the horizontal bold line. The maximum or minimum value within 1.5 x IQR is displayed by the whiskers (vertical lines). The dots are outliers.

### Iron content determination

Between 20 - 30 mg of freeze-dried and ground plant material was dried over night at 62 °C - 65 °C. Dry weight was determined and one milliliter of 67 % trace-metal grade nitric acid was incubated over night at room temperature. Afterwards, samples were heated in a water bath at 80 °C for one hour until all visible particulate matter was completely digested. After cooling down for 10 - 15 min at room temperature, the sample tubes were centrifuged at 4000 rpm for 30 min. 600 µl of the supernatant were collected in a new tube and mixed with 5.4 ml of distilled water. Elemental analysis was performed in the Biocenter-mass spectrometry facility of the University of Cologne by inductively coupled plasma-mass spectrometry (ICP-MS) (Agilent 7700, Agilent Technologies, Santa Clara, United States) using mass standards.

### Histochemical staining of iron with the Perls/3,3′-Diaminobenzidin (DAB) procedure

The histochemical staining with the Perls procedure was conducted with 50 – 100 µm hand-made cross sections according to (Roschzttardtz et al., 2013). The sections were fixed for 1 h using vacuum infiltration in methanol/chloroform/glacial acetic acid (6:3:1). After fixation, plant material was washed three times for 1-2 min with distilled water and incubated under vacuum for 1 h in the pre-warmed (37 °C) stain solution containing 4 % potassium-ferrocyanide (w/v) and 4 % HCl (v/v). After removing the stain solution, sections were washed again three times with distilled water and stored in distilled water at 4 °C either until microscopic analysis or further subjection to the DAB procedure. The intensification using DAB was performed according to (Roschzttardtz et al., 2009). The sections treated with the Perls stain solution (or untreated control sections) were incubated for 1 h in a methanol solution containing 0.01 M NaN_3_ and 0.3 % H_2_O_2_ (v/v).

After removing the solution, plant material was washed three times with 0.1 M phosphate buffer (pH 7.4). For intensification, sections were incubated between 5 and 30 min in 0.1 M phosphate buffer (pH 7.4) containing 0.025 % DAB (w/v), 0.005 % H_2_O_2_ (v/v) and 0.005 % CoCl_2_ (w/v). The reaction was stopped by washing the sections three times for 1-2 min with distilled water. The sections were either stored at 4 °C in distilled water or subjected immediately to microscopic analysis. Microscopy was performed with an Axio Imager 2 microscope (Carl Zeiss Microscopy, Jena, Germany), equipped with an Axiocam 105 color camera (Carl Zeiss Microscopy, Jena, Germany) using bright-field illumination. Images were processed via ZEN 2 (blue edition) software (Carl Zeiss Microscopy, Jena, Germany).

### Ferritin (FER) detection using immunoblot and immunolocalization

Ten milligrams of freeze-dried plant material were used for protein extraction with 2 x SDG buffer (4 % SDS (w/v), 0.2 M DTT, 20 % glycerol (v/v), 0.02 % bromphenol blue). The samples were heated for 10 min at 95 °C and centrifuged for 5 min at 13000 rpm. After centrifugation, 5 µl of each sample was separated in a discontinuous 12 % SDS-polyacrylamide gel. Separated proteins were transferred from the gel to a 0.2 µm nitrocellulose membrane (Amersham Protran 0.2 NC nitrocellulose Western blotting membranes, Cytiva). Rabbit anti-FER IgG (FER antibody) (Agrisera, AS10 674) and rabbit anti-ACTIN (Actin antibody) (Agrisera, AS13 2640) (diluted 1:3000 and 1:5000 in 2.5 % Milk-TBST buffer respectively) were used for immunoblot analysis with a goat anti-rabbit IgG, horseradish peroxidase (HRP)-conjugated (Agrisera, AS09602, diluted 1:10000 in 2.5 % Milk-TBST) as the secondary antibody. Detection of the signals of the enhanced chemiluminescence reaction (Amerscham ECL Select Western Blotting Detection Reagent, Cytiva) was performed via FluorChem Q (proteinsimple). The analysis of the detected FER signals was conducted via AlphaView Software (proteinsimple) with the background-corrected signal normalized to Actin signals.

For the immunolocalization studies, 50 – 100 µm hand-made cross sections were prepared. The immunolocalization analysis of ferritin was performed according to procedures described in (Mohr et al., 2024; Pasternak et al., 2015), whereby sections were first treated with 100 % methanol for 20 min at 37 °C. Sections were incubated with an anti-ferritin antibody, diluted 1:150. Alexa Fluor® 488-conjugated goat anti-rabbit IgG, diluted 1:1000, was applied for detection as the secondary antibody. Microscopy was performed with an Axio Imager 2 microscope (Carl Zeiss Microscopy, Jena, Germany), as described in the previous chapter and in (Mohr et al., 2024).

### Gene expression analysis of iron homeostasis-related genes

The transcriptome data set available under the GEO accession GSE152417 was mined for enriched iron homeostasis terms using topGO (Alexa & Rahnenfuhrer, 2010) with the closest *A. thaliana* ortholog Locus IDs (AGI codes). Publicly available *A. thaliana* gene ontology (GO) annotations (go_ensembl_arabidopsis_thaliana.gaf; downloaded from Gramene) were used. Fisher’s exact test was applied to identify significantly enriched GO terms related to iron homeostasis with p < 0.05 – for list of gene IDs see (Mai et al., 2023) - according to (Sergeeva et al., 2021a). Pearson correlation analysis was conducted via the R software. Correlation coefficients with significant levels of correlation (p < 0.05) are displayed in the diagrams.

## Results

### Iron accumulates in secondary growth tissues of the perennial zone (PZ)

Perennial persistence is restricted to the perennial stem zone (PZ) in *A. alpina*. Since the PZ has secondary growth tissues with concomitant accumulation of carbon-containing polymers, e.g., starch and lipids (Sergeeva et al., 2021a; Sergeeva et al., 2021b), we tested whether iron also accumulated there. We suspected to observe an increase in Fe levels during the development of the PZ. In contrast, we expected that the annual zone (AZ) which is composed of primary growth tissue that does not accumulate C storage compounds in the same manner as the PZ, might not accumulate iron. We analyzed iron accumulation in the plant stem material that we had described previously (Sergeeva et al., 2021a). This material had been collected at different developmental stages from the internodes of lateral stems of *A. alpina* wild-type Paj and of the early flowering mutant *pep1-1*. The growth stages had been assigned to stages I, II, III and II’, depending on the harvesting time point and vernalization growth conditions (details of the growth stages and plant characteristics according to the harvesting scheme represented in Fig. 1A, Fig. 1B). Briefly summarized, stages I and II corresponded to the beginning of secondary growth in the PZ of Paj (stage I before and stage II after vernalization), which then had progressed at stage III. Stage II’ was reached in Paj in the absence of vernalization. The plants had not flowered then, but despite of that had advanced formation of secondary growth tissues in the PZ. At stages II’ and III, plants had distinct stem segments corresponding to a proximal PZ and distal AZ. Secondary growth had developed in a similar manner in *pep1-1* and Paj, while flowering took continuously place in *pep1-1* in contrast to Paj.

When using this material, we found that Fe levels had increased in the PZ internodes of both, Paj and *pep1-1*, at the stages III and II’ versus I and II and in the PZ versus the corresponding AZ segments by roughly two-fold (Fig. 1C). Indeed, Fe contents were highest in the PZ of both investigated Arabis genotypes at stage II’ (35 and 33 µg Fe/g dry matter in Paj and *pep1-1* in the PZ versus 19 and 12 µg/g at stage I, Fig. 1C). Surprisingly, *pep1-1* Fe contents were higher in the AZ regions at stage II’ versus stage III, and at stage III they were also higher in Paj versus *pep1-1*. One explanation for this observation is that Fe accumulated in the AZ to ensure proper flower and silique development taking place at stage II’ in *pep1-1* and stage III in Paj, while stems of *pep1-1* had already senesced at stage III (Fig. 1C). Perhaps, Fe had been exported from the stem segment in the AZ of *pep1-1* at stage III, which might have occurred prior to senescence.

Taken together, these findings point to significantly increased Fe contents in the PZ along a developmental gradient as compared with the AZ. Fe detected in the AZ might be related to the reproductive status of this zone. In addition, similar behavior of Paj and *pep1-1* regarding Fe contents suggests that Fe accumulation in the PZ occurs independent of flowering control by *PEP1* and vernalization.

### Fe is sequestered in ferritin in secondary growth tissues of the PZ

Next, we inspected the localization of Fe more precisely at various stages of the developing PZ and AZ using the Perls Fe staining procedure. The growth stages comprised two early vegetative stages, stage I_PZ and II_PZ without and with initiation of secondary growth (after 4 and 5 weeks), one late vegetative stage, III_PZ with secondary growth (after 7 weeks), and one flowering stage, IV_PZ with advanced secondary growth (after 30 weeks) (Fig. 2A), see also descriptions in (Sergeeva et al., 2021a). For comparison, we investigated the AZ internodes of the 30-week-old flowering stage, either in a region close to the transition of PZ-AZ characterized by short internodes (stage IV_AZ_si) or adjacent to it near the flowers (stage IV_AZ_if) (Fig. 2A), as described (Sergeeva et al., 2021a). We focused the analysis on the *pep1-1* genotype because obtention of respective plant material was well controlled in this mutant.

**Fig. 2.**
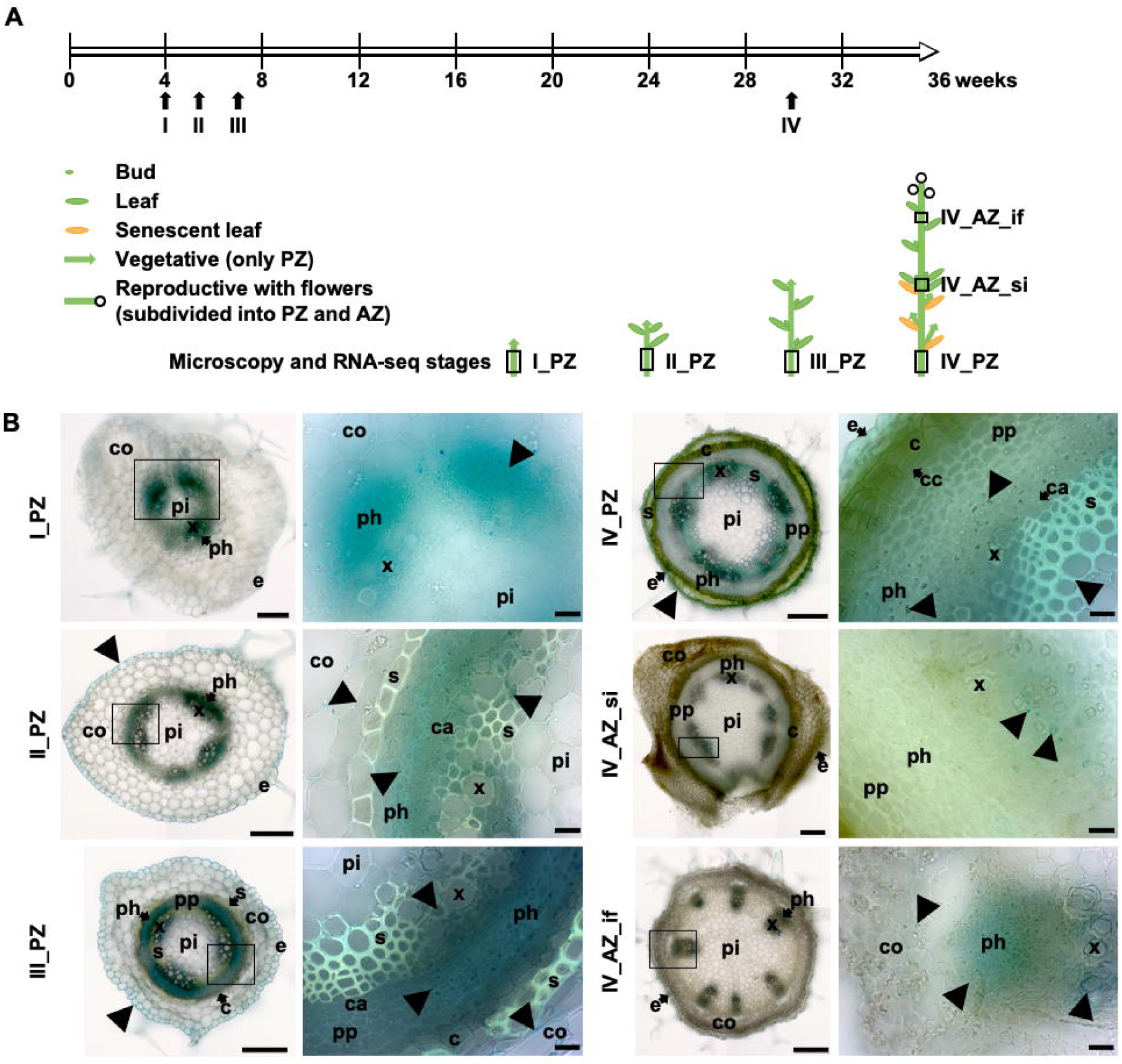
Iron detection in lateral stem internodes of *A. alpina perpetual flowering 1-1* (*pep1-1*) mutant at different developmental stages. (A) Plant growth and harvesting scheme for *A. alpina pep1-1* mutant with schematic representation of the first and second lateral stems architecture originating from lower internodes of the main stem. Open line, long-day conditions at 20 °C; four developmental stages were used for microscopy and RNA-seq analysis, stages I_PZ, II_PZ, III_PZ and IV_PZ as well as IV_AZ_si and IV_AZ_if (Sergeeva et al., 2021a). Lateral stem internodes were dissected into perennial (PZ) and annual (AZ) zones, as indicated. The schematic plant representation was partially adopted from (Lazaro et al., 2018; Vayssières et al., 2020; Wang et al., 2009). (B) Representative lateral stem internode cross sections at stages I_PZ, II_PZ, III_PZ and IV_PZ as well as corresponding to IV_AZ_si and IV_AZ_if. Iron, investigated following Perls staining, in blue, indicated by black triangles. Abbreviations used in microscopic images: c, cork; ca, cambium; co, cortex; e, epidermis; ph, phloem including primary phloem, secondary phloem, phloem parenchyma; pi, pith; pp, secondary phloem parenchyma; s, sclerenchyma; x, xylem including primary xylem, secondary xylem, xylem parenchyma. Arrows point to respective tissues. Scale bars, whole sections, I_PZ, 100 µm; II_PZ, III_PZ, IV_PZ, IV_AZ_si and IV_AZ_if, 200 µm; close-up views, 20 µm.

We detected Fe staining in phloem and xylem at stage I_PZ and II_PZ (Fig. 2B). Stage II_PZ and III_PZ showed presence of Fe also in the apoplast of the epidermis and cortical cells. In addition, Fe was present in a ring of secondary phloem parenchyma at stage III_PZ. At stage IV_PZ, Fe was mainly detectable in the ring of secondary phloem parenchyma of the PZ. Light blue staining of sclerenchyma cells indicated that Fe had accumulated in the cell walls of this tissue. Hence, the secondary phloem which had been previously found associated with storage of carbon (Sergeeva et al., 2021a) was also linked with Fe accumulation. On the other hand, in the short internode region, IV_AZ_si, where secondary growth characteristics, e.g., formation of secondary phloem parenchyma, were present only to some extent, had only single individual blue dots, representing Fe in secondary xylem parenchyma cells. Above, in the AZ, Fe was detectable in phloem and xylem of the IV_AZ_if region (Fig. 2B). Thus, Fe was also accumulating in the vicinity of conductive tissues in the AZ.

Taken together, these observations point to Fe sequestration in the apoplast and in the secondary phloem and xylem parenchyma in the PZ, highlighting a role of the PZ as a perennating storage organ for Fe. Contrary to that, detection of iron in xylem and phloem regions of the AZ and its later disappearance from there might be related to flowering, silique and seed formation and senescence.

It was then interesting to elucidate the nature of Fe sequestration in the secondary phloem of the PZ. Perls staining coupled to diaminobenzidine intensification revealed black dots within plastids present in the secondary phloem and xylem parenchyma of the PZ (Fig. 3A, Fig. 3B). This resembled black ferritin-Fe dots in plastids of Arabidopsis leaves (Roschzttardtz et al., 2013). Ferritin is a protein complex for Fe storage. Ferritin levels correlate with Fe contents, and ferritin detection is used to indicate the presence of Fe accumulation, for example (Kar et al., 2021). We detected ferritin protein in the PZ and AZ (Supplementary Fig. 1). Immunolocalization of ferritin in the first internode of the seven-week-old PZ was aligned with the observed Perls/DAB pattern in secondary phloem and xylem parenchyma (Fig. 3C). Ferritin-associated signal was detected as punctuate labelling in plastids of these tissues (Fig. 3D). Together, these findings indicate that Fe accumulated in ferritin storage complexes in the secondary growth tissues of the PZ of *A. alpina*.

**Fig. 3.**
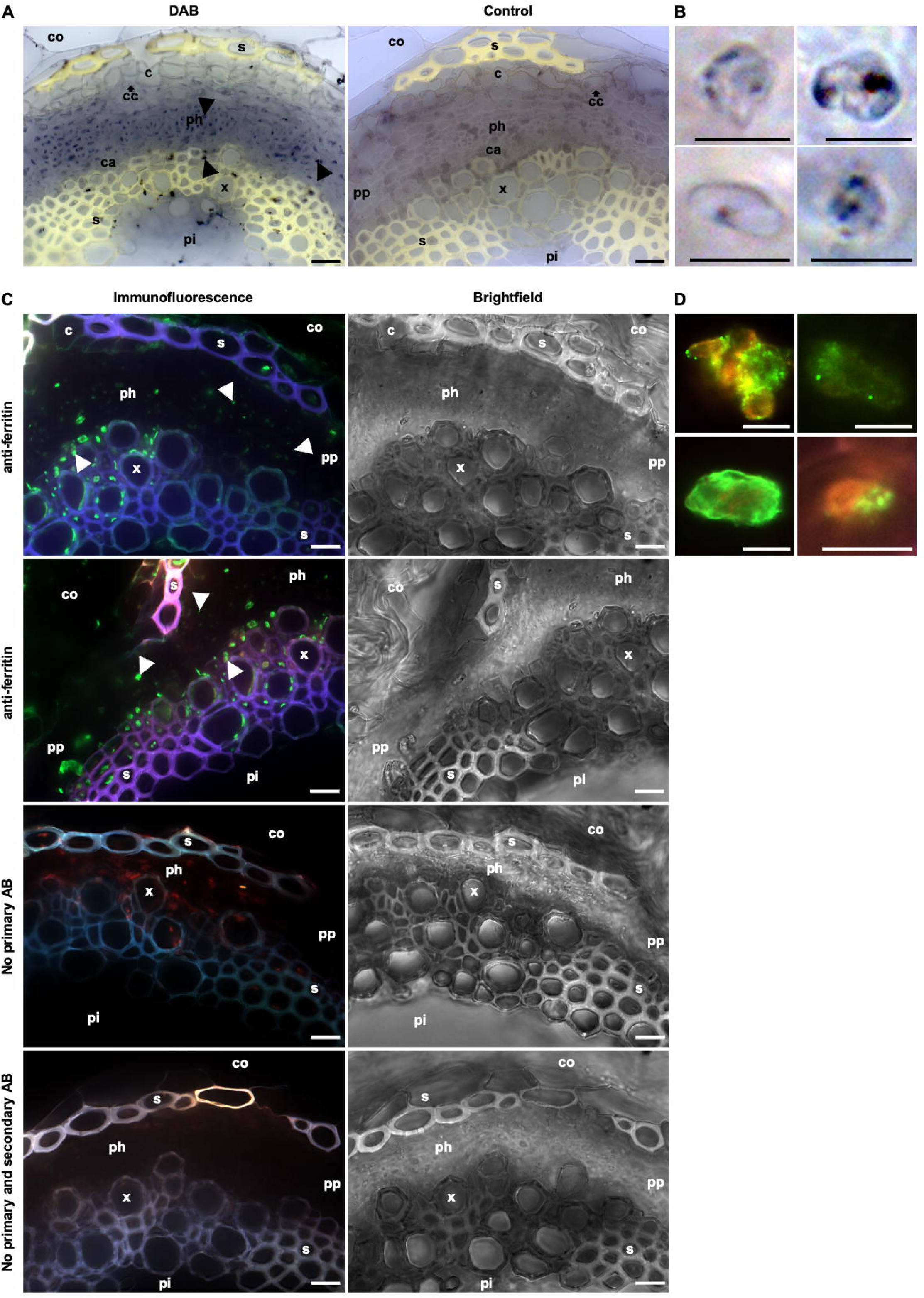
Localization of ferritin in the first internode of the perennial stem zone of *A. alpina perpetual flowering 1-1* (*pep1-1*) mutant. (A) Representative cross sections stained with Perls/DAB, either including Perls reaction (DAB) or not (Control). Iron, stained in black, indicated by black triangles. (B) Magnified images of stained structures detected in xylem and phloem regions represented in (A). (C) Immunolocalization of ferritin. Sections were incubated with an anti-ferritin antibody, Alexa Fluor® 488 conjugated goat anti-rabbit IgG was applied for detection as the secondary antibody. Controls were performed either with secondary antibody only (No primary AB) or without both, primary and secondary antibody (No primary and secondary AB). In addition, grayscale brightfield images of the corresponding regions shown in immunofluorescence pictures are represented. Ferritin localization is displayed in green (white triangles), chlorophyll autofluorescence in red and autofluorescence of lignified cell walls (sclerenchyma, xylem) in blue/purple. (D) Magnified images of ferritin located in chloroplasts detected in xylem and phloem regions represented in (C). Abbreviations used in microscopic images: c, cork; ca, cambium; cc, cork cambium; co, cortex; e, epidermis; ph, phloem including primary phloem, secondary phloem, phloem parenchyma; pi, pith; pp, secondary phloem parenchyma; s, sclerenchyma; x, xylem including primary xylem, secondary xylem, xylem parenchyma. Arrows point to respective tissues. Scale bars, (A), (C), 20 µm; (B), (D), 5 µm.

### Iron homeostasis-related gene expression patterns suggest development-related and tissue-specific differences of Fe homeostasis in the PZ and AZ

To identify which iron homeostasis genes are relevant upon iron accumulation in PZ and AZ, we examined transcriptome data that had been previously investigated from various PZ and AZ growth zones (Sergeeva et al., 2021a). Through many transcriptomic studies performed in the past, a list of Fe homeostasis genes has been gathered for enrichment analysis (Mai et al., 2023).

We used significantly expressed and differentially regulated genes and grouped them according to PZ or AZ stages and subsequently performed GO term enrichment separately for PZ and AZ (Fig. 4A). This resulted in enrichment of two GO terms related to Fe homeostasis, “iron ion transport” and “cellular response to iron ion starvation”. Both GO terms were similarly enriched in PZ and AZ samples, indicating a comparable occurrence and importance of these processes in both zones.

**Fig. 4.**
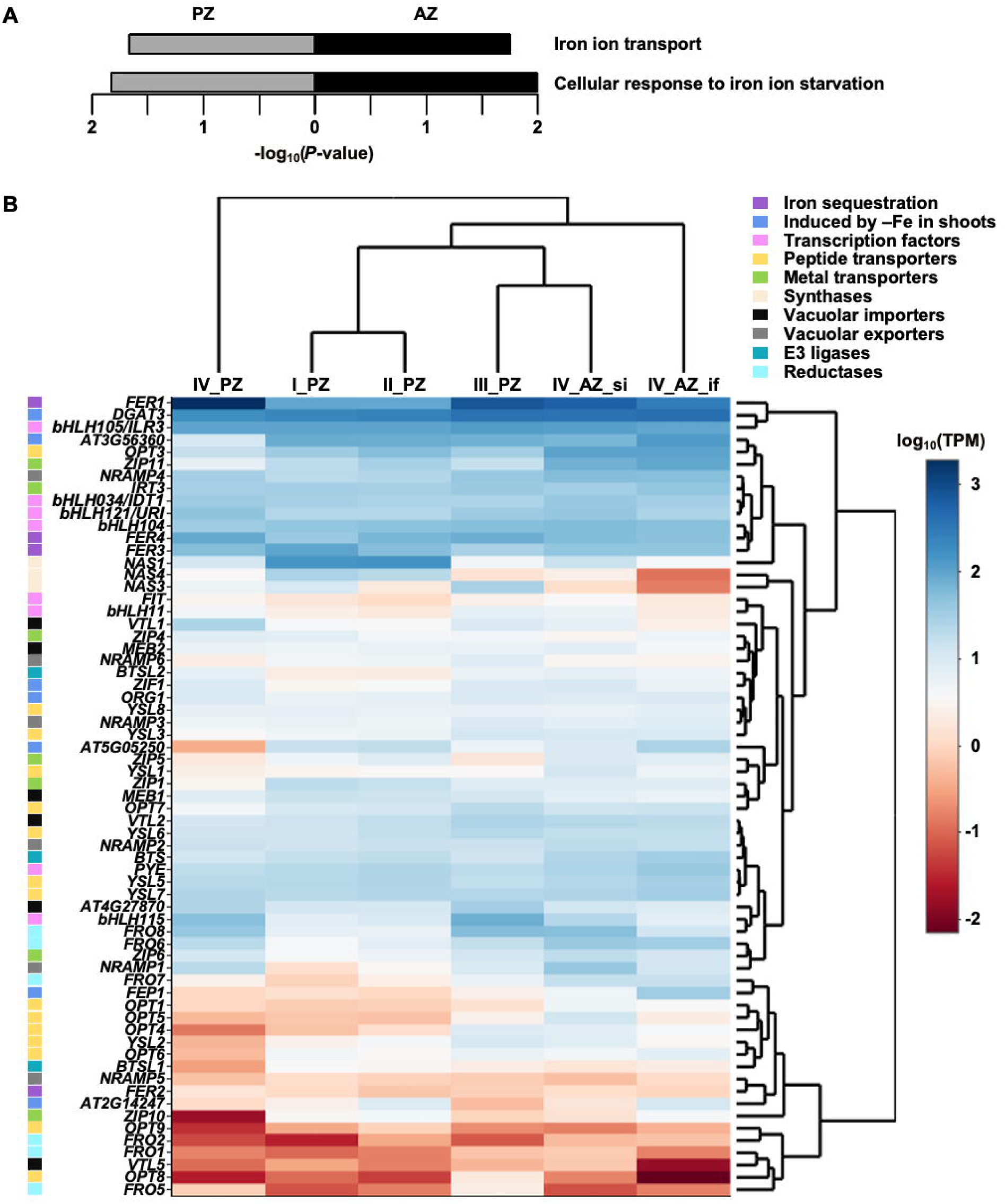
Iron homeostasis-related GO term enrichment analysis of perennial (PZ) versus annual (AZ) lateral stem zones (A) and gene expression profiles of 65 individual genes identified in the RNA-seq data (B) of *A. alpina perpertual flowering 1-1* (*pep1-1*) mutant. (A) PZ represents the combination of stages I_ to IV_PZ (shown in grey), while AZ comprises the sum of stages IV_AZ_si and _if (represented in black). *P*-value of the enriched GO terms < 0.05; *P*-values are represented as –log_10_ values. (B) Gene expression data at stages I_PZ, II_PZ, III_PZ, IV_PZ, IV_AZ_si and IV_AZ_if (stages represented in Fig. 1A, B and Fig. 2A), see also (Sergeeva et al., 2021a) are represented as log_10_ of transcripts per million (TPM) values. Individual genes were assigned to the indicated categories corresponding to iron sequestration (purple), induced by –Fe in shoots (according to Schwarz and Bauer, 2020) (blue), transcription factors (pink), peptide transporters (yellow), metal transporters (green), synthases (beige), vacuolar importers (black), vacuolar exporters (gray), E3 ligases (turquoise) and reductases (light blue). Data are represented as mean (n = 3). Further information about the gene expression data is available in Supplementary Table 1.

We further screened the RNA-seq data for indications about individual gene functions in Fe homeostasis-related processes in *A. alpina* stems (Mai et al., 2023). We identified 65 individual genes and assigned them to ten higher-order categories corresponding to their described functions in various processes of iron acquisition, distribution, storage and regulation (Fig. 4B; Supplementary Table 1), which were “Iron sequestration”, “Induced by -Fe in shoots”, “Transcription factors”, “Peptide transporters”, “Metal transporters”, “Synthases”, “Vacuolar importers”, “Vacuolar exporters”, “E3 ligases” and “Iron reductases”.

The identified Fe homeostasis genes formed four distinct clusters according to their similarity in gene expression levels in the stem internode regions (Fig. 4B, tree above the heat map). One cluster comprised stages I_PZ and II_PZ, grouping closely with the second cluster formed by stages III_PZ and IV_AZ_si. As demonstrated above, these samples all contained cells with secondary growth tissues that had accumulated iron. The third and fourth distinct clusters were stages IV_AZ_if (primary growth characteristics only) and IV_PZ (pronounced secondary growth), whereby the latter was most distant from the other mentioned groups. This suggests that there are development-specific iron homeostasis gene expression patterns.

Clear distinction of individual gene clusters was obtained according to the expression levels of the genes, separating the genes into low-level and high-level expressed genes (Fig. 4B, tree at right side of heat map). Remarkably, the genes assigned to “Iron sequestration”, *FER1*, *FER3* and *FER4*, encoding ferritin proteins, were present among the genes with the highest expression levels. In addition, *FER1* had the highest expression levels at stages III_PZ and IV_PZ, supporting our observation of ferritin-associated iron sequestration in secondary growth tissues of the PZ. Interestingly, we detected expression of several Fe-deficiency-induced genes among the high-level expression genes in the stem segments. These genes correspond to a subset of Fe-deficiency-induced genes in roots and leaves or whole shoots. They are characterized as being strongly induced upon Fe deficiency in a loss of function mutant lacking an essential transcription factor for Fe acquisition in roots named FIT, hence considered “FIT-independent” in Arabidopsis (Schwarz and Bauer, 2020). The FIT-independent genes, and presumably likewise the corresponding *A. alpina* genes, are responsive to iron deficiency cues and involved in internal Fe mobilization and allocation or in regulating the Fe deficiency response cascade, resulting in enhanced Fe acquisition and remobilization capacities under low Fe conditions (Schwarz and Bauer, 2020). In Arabidopsis, the FIT-independent genes are controlled by another level of transcription factors belonging to the subgroup IVc (*bHLH034*, *bHLH104*, *bHLH105*/*ILR3*, *bHLH115*) (Li et al., 2016; Liang et al., 2017; Samira et al., 2018; Tissot et al., 2019; Zhang et al., 2015) and subgroup IVb (*bHLH121*/*URI*, *PYE*, *BHLH011*) (Gao et al., 2019; Kim et al., 2019; Liang et al., 2017; Schwarz and Bauer, 2020; Tanabe et al., 2019). Genes encoding this level of transcription factors were among the very highly expressed genes in the *A. alpina* stems, namely *BHLH105/ILR3*, *BHLH034/IDT1* and *BHLH121/URI*, or among the high-expressed genes, like *BHLH115* and *PYE*. Fine-tuning can occur by the action of E3 ligase BRUTUS (BTS) that can target bHLH IVc factors (Hindt et al., 2017; Long et al., 2010; Selote et al., 2015). The *BTS* gene was also regulated under Fe deficiency and highly expressed in *A. alpina* stems. Among the highly expressed genes were also several genes related to iron mobilization, such as genes encoding vacuolar exporters, like *NRAMP4 (Lanquar et al., 2005)*, and metal chelator-producing nicotianamine synthases (Schuler et al., 2012), and nicotianamine-metal transporter *ZIF1* (Haydon and Cobbett, 2007) which are targets of the above bHLH factors (Long et al., 2010). Hence, the gene expression in stem segments indicates that processes of iron mobilization and export as well as Fe sequestration occur.

On the other hand, among the genes with lowest expression (Fig. 4B) were several iron reductase and oligopeptide transporter genes of the FRO and OPT families (Jain et al., 2014; Mendoza-Cozatl et al., 2014), respectively, and some genes which are associated with Fe deficiency in Arabidopsis roots, such as *FRO2* (Robinson et al., 1999), *BTSL1* (Hindt et al., 2017) and *FEP1/IMA3* (Grillet et al., 2018). BTSL1 is an E3 ligase, counteracted by the small protein FEP1/IMA3 effector (Lichtblau et al. 2022). Presumably, their action is not needed in the PZ and AZ stem segments.

Taken together, the gene expression data indicate that development-related and tissue-specific processes of Fe homeostasis may take place in the stem segments.

Finally, a likely relation between iron levels and gene expression strength in PZ and AZ samples could be uncovered by Pearson correlation analysis (Fig. 5). Interestingly, the genes whose expression levels correlated with Fe levels were clearly distinct in the PZ versus the AZ, and this was the case for all categories of Fe homeostasis genes investigated (Fig. 5). This shows that gene expression differences in the perennial and annual stem parts are correlated either positively or negatively with Fe contents but in different manners, indicating that Fe homeostasis and Fe accumulation take place in the PZ and in the AZ, but are differently controlled and exerted by different genes in these stem segments, suggesting different signaling events.

**Fig. 5.**
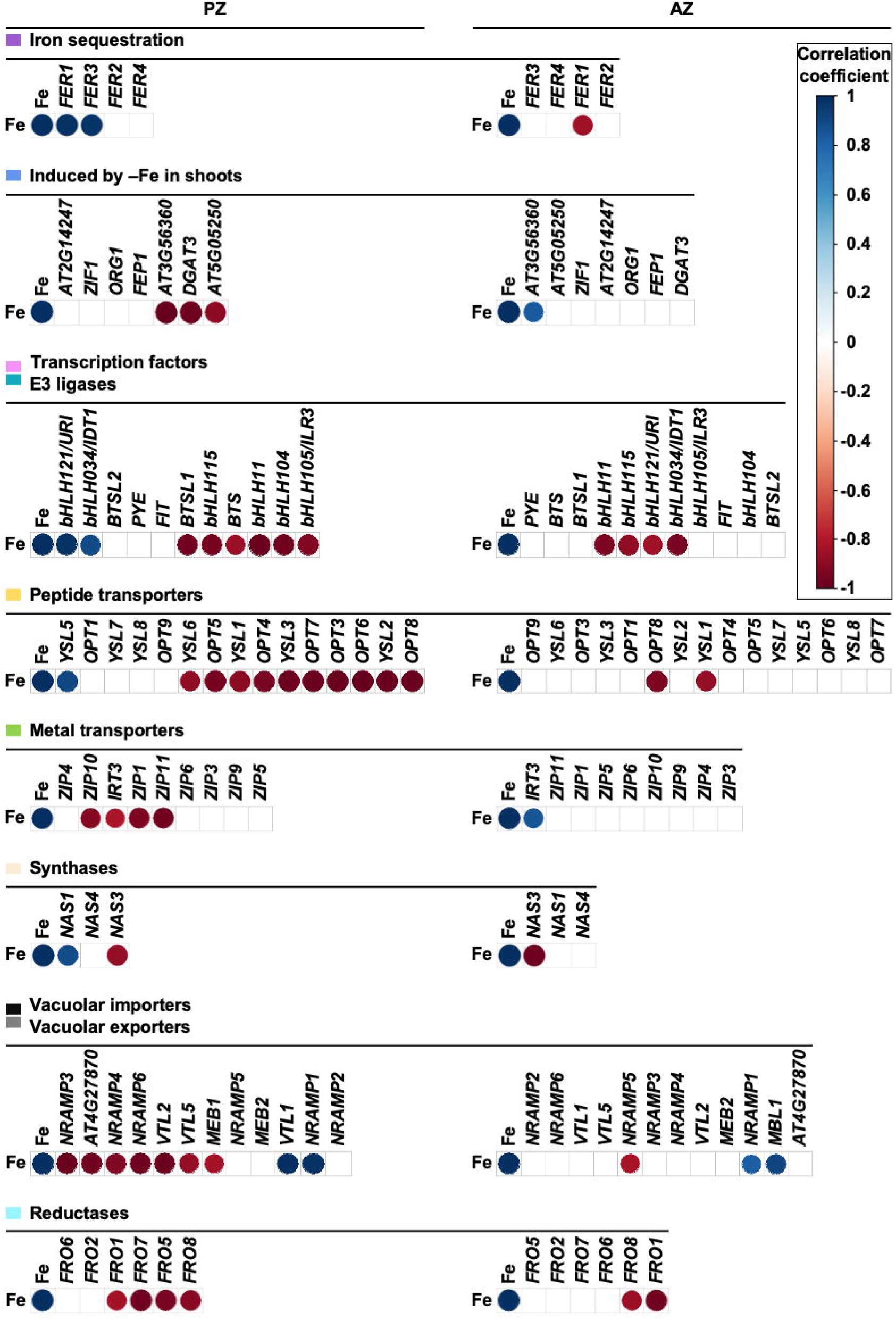
Pearson correlation analysis between gene expression levels of iron homeostasis-related genes/members of the corresponding families and the iron contents of *A. alpina perpetual flowering 1-1* (*pep1-1*) mutant internodes of perennial (PZ) and annual (AZ) lateral stem zones. Stages I, II‘ and III_PZ, IV_PZ were analyzed in the case of the PZ, while the analysis was performed between stage II‘ and IV_AZ_si, _if for AZ (stages represented in Fig. 1A, B and Fig. 2A). Individual genes were assigned to the indicated categories corresponding to iron sequestration (purple), induced by –Fe in shoots (according to (Schwarz and Bauer, 2020)) (blue), transcription factors (pink), peptide transporters (yellow), metal transporters (green), synthases (beige), vacuolar importers (black), vacuolar exporters (gray), E3 ligases (turquoise) and reductases (light blue). Further information about the gene expression data is available in Supplementary Table 1. Correlation coefficients with significant levels of correlation (*P* ≤ 0.05) are represented in the diagrams, while non-significant are not displayed.

## Discussion

Iron (Fe) accumulation in the perennial zone (PZ) of *A. alpina* stems highlights an intricate interplay between plant development and nutrient storage processes. The PZ is a perennating storage region for iron. Fe storage patterns in the PZ show a role for Fe accumulation in secondary growth tissues. This underscores the PZ’s importance in storing essential nutrients to support subsequent growth seasons and it shows the capacity of secondary growth tissues in stems to act as sinks for Fe.

The observed accumulation of iron like of other nutrients in the PZ mirrors findings in other perennial plants where nutrient storage in woody tissues is critical for overwintering and regrowth. Studies on woody perennials like Populus trees have similarly highlighted the importance of secondary growth tissues in nutrient storage (Cooke and Weih, 2005; Wildhagen et al., 2010) and presumably iron can also be stored there.

The increase in Fe levels during the transition from primary to secondary growth in the PZ suggests a highly regulated mechanism of nutrient remobilization and storage. This aligns with previous work showing that storage tissues in perennials accumulate and redistribute nutrients depending on developmental cues (Zapata et al., 2004). The increased Fe content in the PZ supports the hypothesis that secondary growth tissues are optimized for long-term storage and remobilization, critical for perennial plants undergoing seasonal growth cycles.

The developmental gradient of Fe accumulation indicates that storage organ characteristics are not only spatially but also temporally regulated. This temporal regulation may ensure that Fe is available in periods of high demand, such as during flowering, fruit development and seed maturation. For example, there is a lower amount of iron in the *pep1-1* AZ versus PZ at Stage III compared to Paj. Because the Fe content is roughly equal in *pep1-1* and Paj at Stage II, this may indicate a more pronounced or earlier onset of Fe re-localization from AZ to PZ in *pep1-1*. The observation that both lines appear to have more Fe at Stage III than at Stage II, could indicate that this occurs along with increased Fe uptake during or right before Stage III or due to decreased growth, biomass production and/or Fe demand at Stage III. The findings that Fe accumulation occurs independent of vernalization and PEP1-mediated flowering control suggest a robust underlying regulatory system that could be advantageous in varying growing environments and conditions.

The distinct gene expression patterns related to iron homeostasis further reveal development-specific adaptations. Negative correlations between gene expression and Fe levels, e.g. low Fe and high gene expression, may indicate that genes control internal mobilization and redistribution rather than storage of Fe in the same plant segments, while positive correlation, e.g. high Fe levels and high gene expression could be a sign of an involvement of genes in Fe storage.

The high expression of ferritin and other iron sequestration genes in the PZ, where Fe accumulates, aligns with the physiological need to store Fe securely in response to iron availability and developmental signals. The role of ferritin in Fe sequestration observed in this study also parallels findings in Arabidopsis leaves, where ferritin serves as a major iron storage molecule, protecting plants from oxidative stress caused by free iron (Roschzttardtz et al., 2013). Among the genes related to Fe sequestration, gene expression levels of *FER1* and *FER3* positively correlated with the Fe levels in the PZ, but this was not the case in the AZ, indicating that an increase of Fe in the PZ is correlated with ferritin formation due to FER1 and FER3.

The presence of Fe-deficiency induced genes among the high-expressed genes in the PZ suggests a complex regulatory network that adjusts Fe homeostasis in response to internal and environmental cues, resembling mechanisms known in Arabidopsis where FIT-independent pathways play a vital role in systemic Fe regulation (Schwarz and Bauer, 2020).

The genes involved in the regulation of Fe homeostasis, such as transcription factors and E3 ligases, showed positive and negative correlations with Fe levels. While positive correlations were identified between *bHLH121* and *bHLH034* and Fe levels in the PZ, *bHLH115*, *bHLH104*, *bHLH105*, *BTS* and *BTSL1* were negatively correlated with the quantified Fe present in the PZ tissues. In the AZ, negative correlations included *bHLH115*, *bHLH121* and *bHLH034*. Perhaps different mechanisms and signals control Fe transport in AZ and in PZ. Moreover, the findings indicate a specialization among the bHLH TFs for controlling processes in the AZ and PZ, in agreement with differing phenotypes of Fe deficiency-related *bhlh* mutants in Arabidopsis (Gao et al., 2024).

YSL transporters, such as AtYSL1, AtYSL2 and AtYSL3, are involved in Arabidopsis in Fe distribution in plant tissues (Waters et al., 2006). In addition, OPT3 plays a role in Fe transport in the phloem and in mediation of Fe shoot to root signaling (Mendoza-Cozatl et al., 2014). Here, we analyzed the whole set of identified genes of these families and found positive correlations between *YSL5* expression and Fe levels in the PZ and negative correlations regarding *YSL6*, *OPT5*, *YSL1*, *OPT4*, *YSL3*, *OPT7*, *OPT3*, *OPT6*, *YSL2* and *OPT8*. Two negative correlations were detected in the AZ, which included *OPT8* and *YSL1*.

Transport of metal ions in plant tissues involves activity of ZIP transporters (Guerinot, 2000). We identified several negative and positive correlations between the gene expression levels of these metal transporters and the Fe levels in Arabis tissues. Negative correlations were present in the PZ and included *ZIP10*, *IRT3*, *ZIP1* and *ZIP11*. One positive correlation between *IRT3* expression and quantified Fe amounts was detected in the AZ.

Nicotianamine synthases, such as NAS4, generate the metal chelator nicotianamine, which is involved in Fe allocation to young sink tissues (Schuler et al., 2012). Our analysis revealed one positive correlation between *NAS1* gene expression levels and the quantified Fe amounts in the PZ. One negative correlation was detected regarding *NAS3* in the PZ and in the AZ. This specialization of NAS genes in different aspects of iron homeostasis is not unexpected as it is also found in *A. thaliana* (Klatte et al., 2009; Schuler et al., 2011; Schuler et al., 2012). However, the gene functions may differ between Arabidopsis and Arabis, e.g. in *A. thaliana*, *NAS3* expression is more pronounced under sufficient than deficient Fe supply in seedlings (Klatte et al., 2009).

Among the genes related to vacuolar metal importers and exporters (Kim et al., 2006; Lanquar et al., 2005), *VTL1* and *NRAMP1* positively correlated with the quantified Fe levels in the PZ. Negative correlations in the PZ were detected for *NRAMP3*, *NRAMP4*, *NRAMP6*, *VTL2*, *VTL5* and *MEB1*. In the AZ, positive correlations included *NRAMP1* and *MBL1*, while one negative correlation was detected for *NRAMP5* here.

Finally, among the identified family members of ferric chelate reductases, only negative correlations between the gene expression levels of these genes and the quantified Fe amounts were detected in the PZ and in the AZ of *A. alpina*. Such correlations in the PZ included *FRO1*, *FRO7*, *FRO5* and *FRO8*, while in the AZ, these were *FRO8* and *FRO1*.

Future research should explore the signaling pathways that govern Fe accumulation and gene regulation in perennial tissues, potentially involving cross-talk between iron homeostasis and carbohydrate metabolism pathways. Investigating the specific roles of identified high-expression Fe-regulatory genes in the PZ and comparing their functions across different perennial species could provide deeper insights into the evolutionary adaptations of nutrient storage mechanisms in perennials. Previous investigation in *A. alpina* highlighted a role of cytokinins in carbon sequestration and differentiation of the PZ (Sergeeva et al., 2021a). Likewise cytokinins might act to control the Fe homeostasis processes in there.

## Conclusions and perspectives

In conclusion, this study accentuates the essential role of the PZ in nutrient storage and mobilization in *A. alpina*, demonstrating a sophisticated balance of iron sequestration that supports perennial growth cycles. Understanding these mechanisms not only enhances our knowledge of plant physiology but also has implications for improving nutrient management in crops, highlighting the potential for bioengineering plants with optimized nutrient storage capacities.

## Abbreviations

AZ: annual zone
DM: dry matter
GO: gene ontology
if: inflorescence
Paj: Pajares
*pep1-1*: *perpetual flowering 1-1*
PZ: perennial zone
si: short internode

## Acknowledgements

This work received funding by Deutsche Forschungsgemeinschaft (DFG, German Research Foundation) under Germanýs Excellence Strategy – EXC-2048/1 – project ID 390686111. The authors thank the University of Cologne Biocenter-MS platform for ICP-MS analysis. We acknowledge the assistance of AI App, based on the GPT-4 model developed by OpenAI, for language correction, style, summarizing and text shortening.

## Author contributions

Conceptualization: A.S., P.B.; Formal analysis: A.S., H.J.M.; Investigation: A.S., H.J.M.; Writing -original draft: A.S., P.B.; Writing -review & editing: A.S., P.B., H.J.M.; Visualization: A.S.; Supervision: P.B.; Funding acquisition: P.B.

---

**Supplementary Fig. 1:** Immunodetection of ferritin in lateral stem internodes of *A. alpina* Pajares (Paj, wild type) and its *perpetual flowering 1-1* (*pep1-1*) mutant derivative at different developmental stages. Three developmental stages were used for immunodetection analysis, stages I, III and II‘ (see also Sergeeva *et al*. 2021 and Fig. 1B). Lateral stems were subdivided into perennial (PZ) and annual (AZ) zones. Rabbit anti-FER IgG (FER) and rabbit anti-ACTIN (Actin) (diluted 1:3000 and 1:5000, respectively) were used for immunoblot analysis with a goat anti-rabbit IgG, HRP conjugated (diluted 1:10000) as the secondary antibody. The analysis of the detected FER signals was conducted with the background-corrected signal normalization to Actin immunostaining.

**Supplementary Table 1:** RNA-seq gene expression data of selected iron homeostasis-related genes represented in Fig. 4B and Fig. 5.

