## Supplementary Fig. 1 for "Iron homeostasis in the annual and perennial stem zones of *Arabis alpina*"

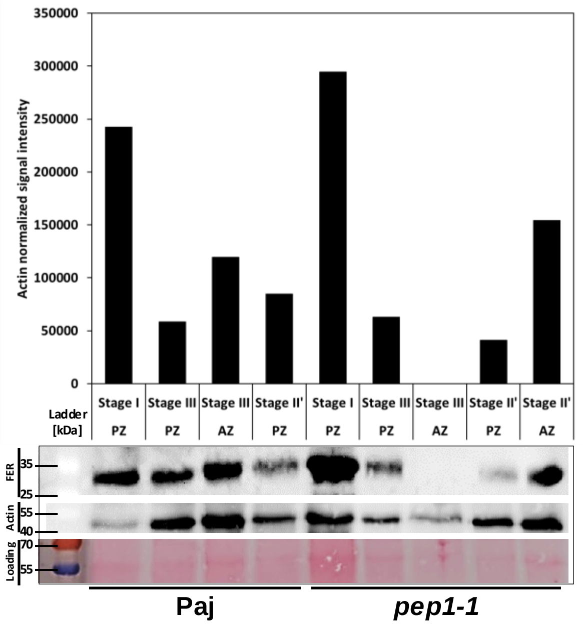


**Supplementary Fig. 1.** Immunodetection of ferritin in lateral stem internodes of *A. alpina* Pajares (Paj, wild type) and its *perpetual flowering 1-1* (*pep1-1*) mutant derivative at different developmental stages. Three developmental stages were used for immunodetection analysis, stages I, III and II‘ (see also Sergeeva *et al.* 2021 and Fig. 1B). Lateral stems were subdivided into perennial (PZ) and annual (AZ) zones. Rabbit anti-FER IgG (FER) and rabbit anti-ACTIN (Actin) (diluted 1:3000 and 1:5000, respectively) were used for immunoblot analysis with a goat anti-rabbit IgG, HRP conjugated (diluted 1:10000) as the secondary antibody. The analysis of the detected FER signals was conducted with the background-corrected signal normalization to Actin immunostaining.
